# High habitat-specificity in fungal communities of an oligo-mesotrophic, temperate lake

**DOI:** 10.1101/056929

**Authors:** Christian Wurzbacher, Norman Warthmann, Elizabeth Bourne, Katrin Attermeyer, Martin Allgaier, Jeff R. Powell, Harald Detering, Susan Mbedi, Hans-Peter Grossart, Michael T. Monaghan

## Abstract

Freshwater fungi are a poorly studied paraphyletic group that include a high diversity of phyla. Most studies of aquatic fungal diversity have focussed on single habitats, thus the linkage between habitat heterogeneity and fungal diversity remains largely unexplored. We took 216 samples from 54 locations representing eight different habitats in meso-oligotrophic, temperate Lake Stechlin in northern Germany, including the pelagic and littoral water column, sediments, and biotic substrates. We pyrosequenced with an universal eukaryotic marker within the ribosomal large subunit (LSU) in order to compare fungal diversity, community structure, and species turnover among habitats. Our analysis recovered 1024 fungal OTUs (97% criterion). Diversity was highest in the sediment, biofilms, and benthic samples (293-428 OTUs), intermediate in water and reed samples (36-64 OTUs), and lowest in plankton (8 OTUs) samples. NMDS clustering clearly grouped the eight studied habitats into six clusters, indicating that total diversity was strongly influenced by turnover among habitats. Fungal communities exhibited pronounced changes at the levels of phylum and order along a gradient from littoral to pelagic habitats. The large majority of OTUs could not be classified below the order level due to the lack of aquatic fungal entries in taxonomic databases. Our study provides a first estimate of lake-wide fungal diversity and highlights the important contribution of habitat-specificity to total fungal diversity. This remarkable diversity is probably an underestimate, because most lakes undergo seasonal changes and previous studies have uncovered differences in fungal communities among lakes.

## Introduction

Aquatic fungi play an important role in the cycling of carbon and nutrients (Gleason et al. 2008; Wurzbacher et al. 2010; Jobard et al. 2010; Grossart and Rojas-Jimenez 2016). The degradation of recalcitrant plant and animal residues is carried out by a number of poorly known groups within the phyla Chytridiomycota and Rozellomycota, yeasts, and hyphomycete lineages of Ascomycetes (reviewed by Wurzbacher et al. 2010; Jobard et al. 2010). Parasitism by Chytridiomycota facilitates the trophic transfer of nutrients from otherwise inedible phytoplankton to filter-feeding zooplankton (termed the “mycoloop”; Kagami et al. 2014, 2007). Aquatic fungi can also be symbionts, for example endophytic or mycorrhiza-forming fungi (Kohout et al. 2012). Despite their importance to ecological function and their diversity in nutrition modes, the biodiversity of aquatic fungi remains poorly known.

Estimates of total fungal diversity range from 0.5 - 10 M species worldwide (Blackwell 2011, Bass and Richards 2011). Of these, roughly 100,000 species are described, only ca. 3000 from aquatic habitats (Shearer et al. 2007; Tsui et al. 2016). The low diversity of aquatic fungal species compared to terrestrial (e.g., soil) ecosystems partly results from the fact that mycological studies in aquatic systems remain rare. Apart from a few well studied lentic ecosystems and wetlands (Wong et al. 1998; Shearer et al. 2007; Gulis et al. 2009; Krauss et al. 2011), the total diversity of aquatic fungi has not been addressed in terms of the link of habitat diversity to fungal diversity. Most studies in freshwaters have focussed on the open water, leaf litter or emergent macrophytes (e.g., *Typha, Phragmites)* with a strong focus on marshlands (reviewed in Kuehn 2008). Studies in the water column have often concentrated on seasonal patterns (e.g., van Donk and Ringelberg 1983; Holfeld 1998; Lefèvre et al. 2012; Rasconi et al. 2012) or on the comparison of different lakes (e.g. Zhao et al. 2011; Lefèvre et al. 2012; Taib et al. 2013). Several studies have found evidence for vertical and horizontal structuring of fungal communities in the water column (Lefèvre et al. 2007; Chen et al. 2008; Lepère et al. 2010), suggesting that there is an important spatial component of diversity. A recent meta-analysis found that aquatic fungi clustered in habitat-specific biomes, with freshwater biomes having the highest fungal diversity at the phylum level (Panzer et al. 2015). The authors attributed this to the high substrate diversity and temporal dynamics of environmental parameters in freshwater systems.

Considering the multitude of available niches and fungal lifestyles in aquatic habitats (Wurzbacher et al. 2010), the actual species number of aquatic fungi is likely to be much higher than what is currently recognized. Freshwater systems contain a large amount of habitat diversity including the boundaries that connect them to terrestrial and groundwater ecosystems (Vadeboncoeur et al. 2002; Schindler and Scheuerell 2002). Temperate, stratified lakes encompass horizontal gradients such as from shallow (littoral zone) to open water (pelagic zone) habitats, as well as vertical gradients from the epilimnion (often euphotic) to hypolimnion (often aphotic) to the aphotic sediment. Shore regions are transition zones between terrestrial and aquatic habitats and include macrostructures such as aquatic macrophytes, animals, plant debris, and biofilms. They may thus be “hot spots” of aquatic, amphibious, and terrestrial fungi (Wurzbacher et al. 2010). In contrast, pelagic habitats have few macrostructures, and fungi may be limited to planktonic substrates such as dissolved organic matter (DOM) and organisms such as phyto- and zooplankton (living or dead). In particular, accompanying the change of substrate from coarse particulate organic matter (CPOM) near the edges of the lake to fine particulate organic matter (FPOM) in the open water, Dikarya are expected to be replaced by less abundant yeasts and Chytridiomycota (Wurzbacher et al. 2010). We hypothesize that this change in “fungal morphotypes” towards unicellular fungi depends on the change in the amount and size of available substrates present in the various lake habitats.

We examined fungal diversity in a temperate lake in North-East Germany (Lake Stechlin) in order to examine the effect of habitat specificity on fungal diversity, and to test the morphotype hypothesis of the littoral-pelagic gradient. We expected changes in diversity and composition in the fungal community to be driven by habitat variability, specifically that the diversity in substrate size and structure (e.g., zooplankton, macrophytes) would support a heterogeneous fungal community. A total of eight representative habitat types at 54 sampling stations were sampled at three time points in spring and analyzed using pyrosequencing of the large ribosomal subunit (LSU) as a universal eukaryotic marker. Our findings revealed a surprisingly high fungal diversity with a high taxonomic turnover among water, plankton, sediment, biofilm, and macrophyte habitats.

## Methods

### Sampling site

Lake Stechlin is a deep (max. depth: 69.5 m), oligo-mesotrophic, dimictic hard-water lake in northern Germany (53° 10’ N; 13° 02’ E). It has a surface area of 4.25 km^2^ and is divided into three distinct basins (Figure 1). There is a littoral reed belt (*Phragmites australis*) that is frequently interrupted by areas of underwater macrophytes (mainly *Characea* and *Potamogeton*) and the lake is surrounded by mixed forest dominated by *Pinus sylvestris* and *Fagus sylvatica.* Lake Stechlin is part of the global lake ecological observatory network (GLEON) and has been monitored since 1959 (Casper 1985). During the course of our study (April - June 2010), the phytoplankton community was dominated by diatoms and filamentous cyanobacteria (*Dolichospermum flos-aquae*) and the nutrient status of the lake during the sampling period is detailed in the Online Resource 1.

**Figure 1.**
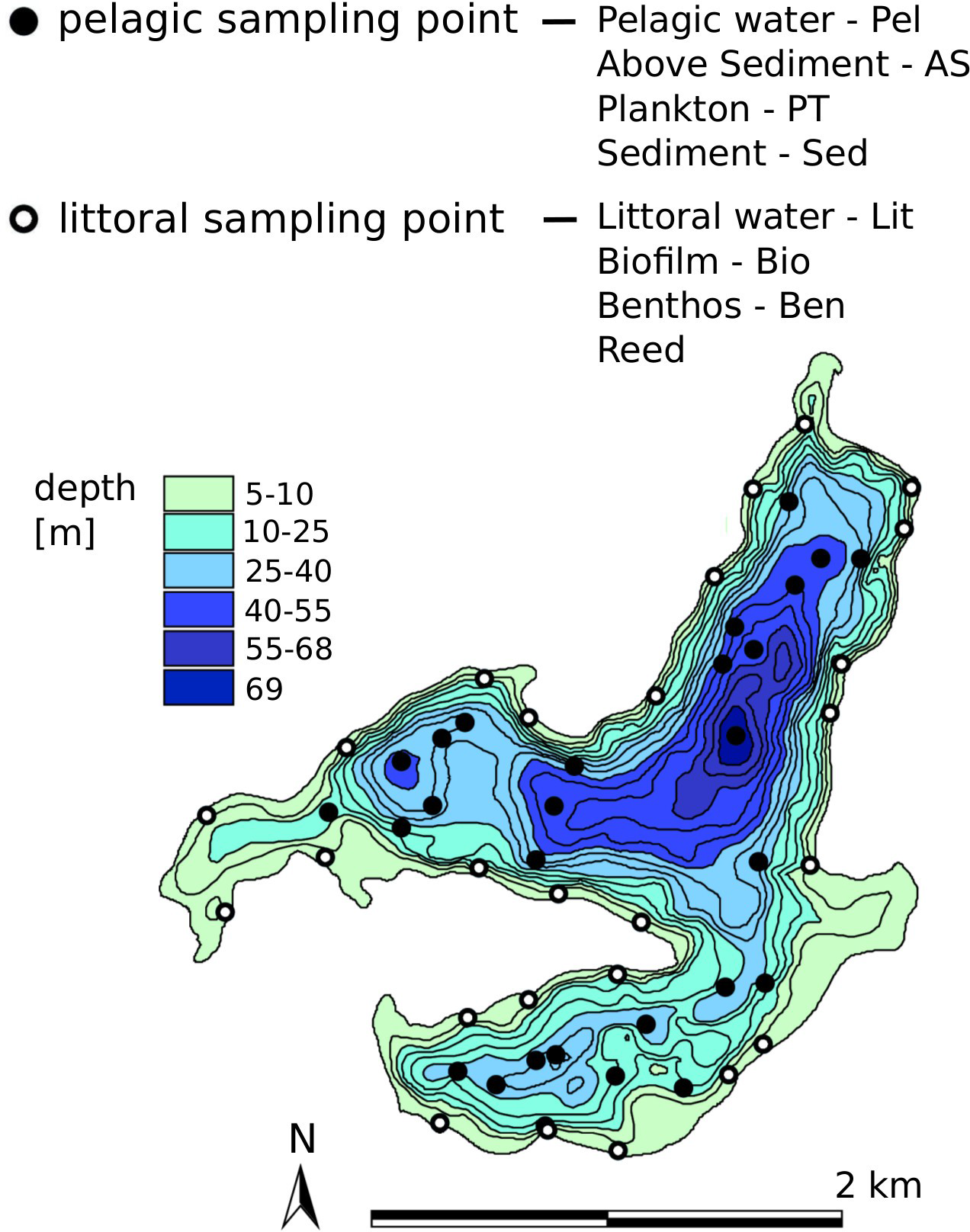
Sampling sites in Lake Stechlin. Integrated water samples, above-sediment water, plankton (> 55 µm), and sediment were taken from pelagic locations. Surface water samples, reed plants (Phragmites australis), biofilm samples (from stone, wood and macrophytes) and benthic samples (detritus, macrozoobenthos) were taken from littoral locations.

### Sampling

We sampled eight different habitat types (Table 1) at three time points encompassing spring and early summer 2010 (8-9 April; 11-12 May; and 9-10 June). Sampling was carried out relatively early in the year to avoid an over-representation of wood-degrading Basidiomycetes that are introduced from the surrounding forest, most of which release spores from July to November (authors, pers. obs.). Our sampling was designed to represent both pelagic (defined here as > 100 m from shore and in areas with >20 m depth) and littoral (<10 m from shore) habitats. Some locations contained multiple habitats and thus comprised a total of 54 sampling locations (Figure 1). Habitats were defined as follows: “Pelagic” samples consisted of a 1 l water sample integrated from 1 m below the surface, at mid-depth, and at 2-3 m above the sediment using a Niskin-type water sampler (Hydro-Bios, Germany); “Plankton” was obtained from an integrated sample (surface to 2-3 m above sediment) from a plankton net (55 µm mesh; Hydro-Bios, Germany); “Above Sediment” was a water sample from 0-20 cm above the sediment that was retrieved together with “Sediment” -1 ml of the uppermost cm of the core-using a sediment corer (6 cm diameter; Uwitec, Austria). “Littoral” samples consisted of a 1-L water sample taken from 0.5-1 m depth in the littoral zone; “Reed” samples were taken from aerial, submerged, and rhizosphere parts of reed plants, following the physical removal of biofilm; “Biofilm” samples were taken from stones, woody debris, and reed stems (removed using a scalpel); and “Benthos” consisted of detritus and zoobenthos sampled from the littoral zone using a sediment grabber. Each of the eight habitat types was sampled at 3 locations in each of the 3 basins at each of the 3 time points (n = 27), for a total of 216 samples. Samples were pooled by combining one sample from each of the three basins, resulting in 3 representative samples of each habitat per time point. These were further pooled for sequencing analysis (see below). Water samples were filtered on a 0.22 µm Sterivex filter (Millipore), plankton-net samples were filtered onto a 12 µm cellulose acetate filter (Sartorius), and 1 ml of sediment was transferred to a cryotube for storage. All samples and filters were stored at -80°C until further processing.

**Table 1.** Total number of sequencing reads analysed and total number of OTUs recovered using alignment-based clustering (97% criterion) for each habitat in Lake Stechlin. Habitat abbreviations used in Fig 2 and Fig. 3 are indicated to the right of habitat names. Reported values were obtained from analysis of 3 samples (1 sample from reed habitat), each of which consisted of pooled DNA from three time points (April-June 2010). Values in parentheses indicate the percentage of total reads and total OTUs that were classified as fungi.

| Habitat | n | Total reads | Fungal reads | Total OTUs | Fungal OTUs | Chao (SE) |
| --- | --- | --- | --- | --- | --- | --- |
| *Pelagic* |  |  |  |  |  |  |
| Pelagic water - Pel | 3 | 19795 | 219 (1%) | 470 | 36 (8%) | 490.5 (49.3) |
| Above sediment -AS | 3 | 18701 | 343 (2%) | 592 | 62 (11%) | 616.2 (60.0) |
| Plankton - PT | 3 | 17902 | 85 (<1%) | 165 | 8 (5%) | 185.5 (29.7) |
| Sediment - Sed | 3 | 79190 | 4011 (5%) | 1753 | 428 (24%) | 908.1 (75.7) |
| *Littoral* |  |  |  |  |  |  |
| Littoral water - Lit | 3 | 29128 | 215 (<1%) | 521 | 40 (8%) | 361.5 (35.4) |
| Biofilm - Bio | 3 | 26302 | 2668 (10%) | 1049 | 293 (28%) | 1104.9 (87.5) |
| Benthos - Ben | 3 | 59169 | 42836 (72%) | 1069 | 424 (40%) | 1055.4 (98.2) |
| Reed | 1 | 4277 | 3760 (89%) | 177 | 64 (36%) | -- |

**Figure 2.**
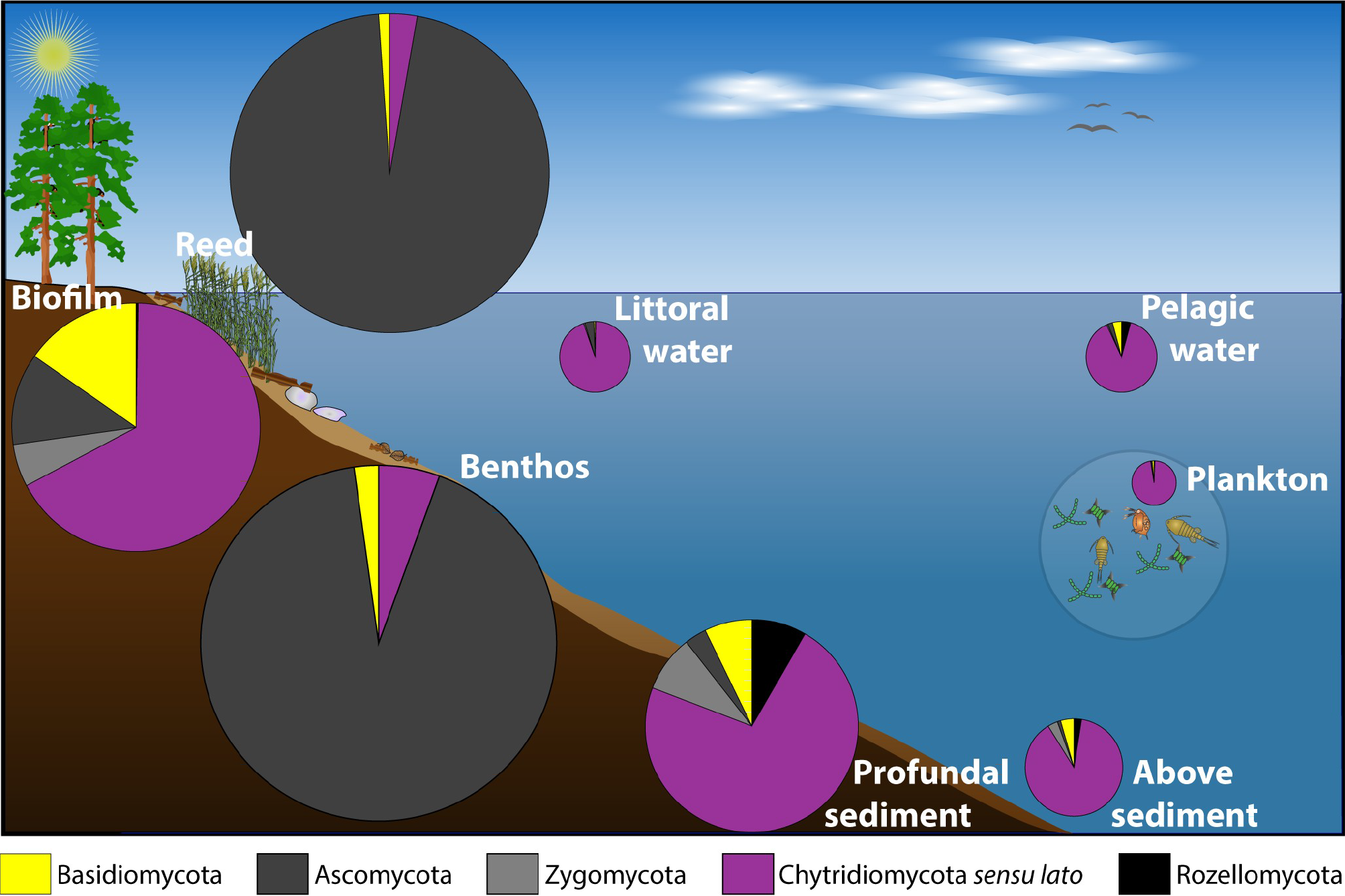
Contribution of fungal phyla within each of the lakes habitat. Pie charts are scaled by the contribution of fungal diversity to the total eukaryotic diversity in each habitat, taken from Table 1 (range: 5-40%).

**Figure 3.**
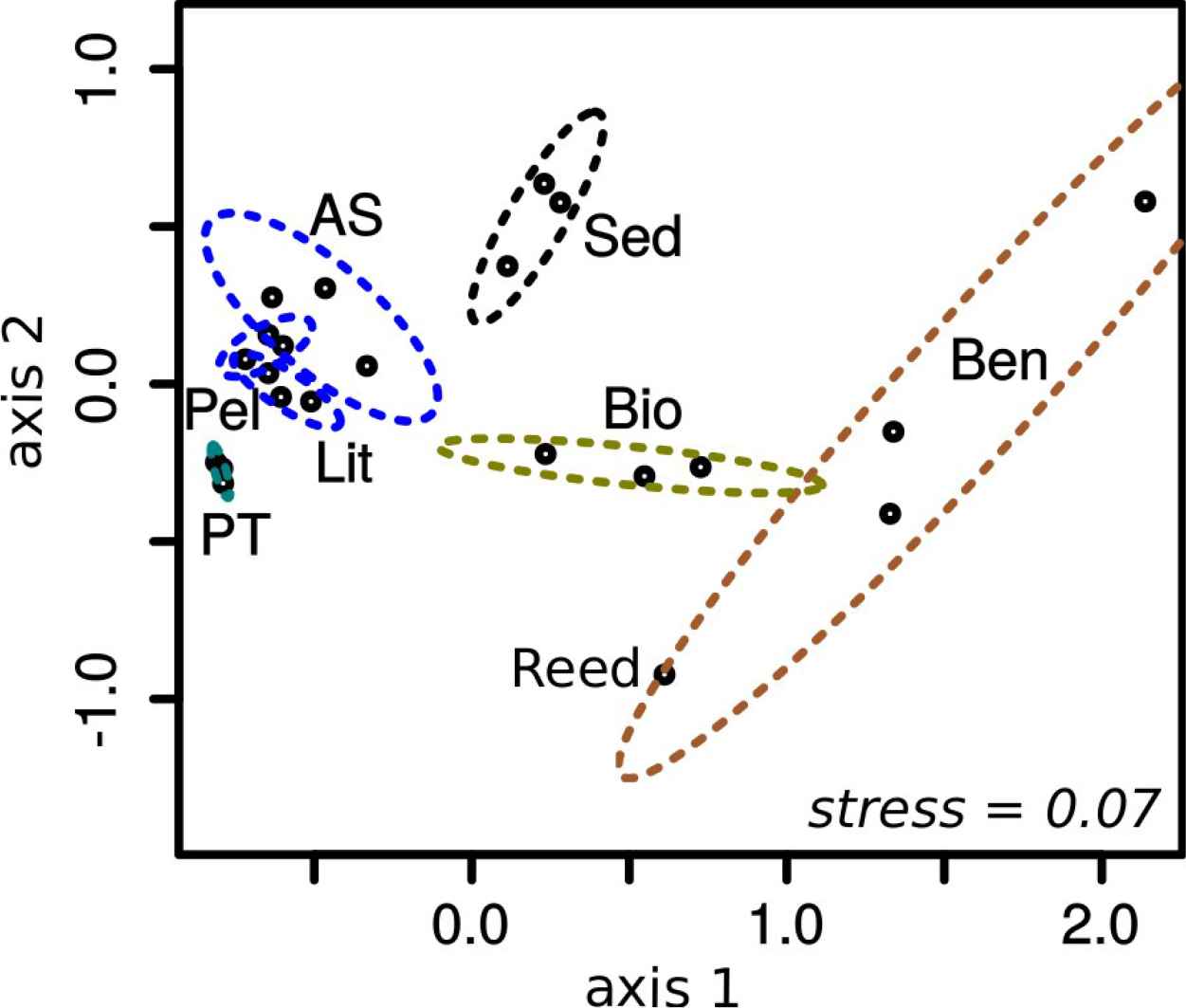
Habitat specificity of the fungal community in Lake Stechlin. NMDS plots based on the fungal OTU matrix (1042 OTUs). Ordination is based on Cao distances (Cao et al. 1997), which are insensitive to differences in sampling effort. Ellipses are based on two standard deviations around habitat centroids based on a confidence levelof 0.95. Habitat codes are taken from Table 1.

### DNA extraction

Total DNA was extracted using the Power Soil kit (MoBio Laboratories, Carlsbad, USA) for Sediment samples; the Qiagen Plant kit (Qiagen, Hilden, Germany) for Reed, Biofilm, and Benthos samples; and using the Qiagen Blood & Tissue kit for Littoral, Pelagic, and Above Sediment water samples, as well as for Plankton samples. Manufacturers’ instructions were followed with the following modifications: Reed and Benthos samples were homogenized with a mill (Pulverisette 9, rpm=self-optimize speed, 20 sec, Fritsch, Germany) and all other samples were subjected to a bead-beating step prior to extraction (MMX400, 2×2 min, f=30 sec^-1^, Retsch, Germany). We added 20 µl Proteinase K (Qiagen, Netherlands) to the lysis buffer for Sediment, Reed, Biofilm, and Benthos samples, and incubated these for 1 h at 56°C. DNA concentrations were measured using a PicoGreen assay (Invitrogen, USA) whereby ~ 20 ng of DNA was used as template for PCR.

### Library preparation for pyrosequencing

All samples were analysed for the D1/D2 variable region of the LSU with the eukaryotic primers NLF184cw (TACCCGCTGAAYTTAAGCATAT; modified from Van der Auwera et al. 1994) and Euk573rev (AGACTCCTTGGTCCRTGT; modified from NLR818, Van der Auwera et al. 1994). After *in silico* tests using TestPrime (Klindworth et al. 2012) we found the primer pair covered 84% of all eukaryotes deposited in the SILVA database (LSU r123 version) when allowing for two mismatches, with none in the last 3 bp of the 3’ region. It potentially excludes single eukaryotic lineages within Amoebozoa, Excavata, Cercozoa. Peronosporomycetes (oomycetese were covered at 76%. Within fungi it covers 93.4% of deposited sequences in all phyla, except Microsporidia. Among the fungal phyla, the lowest coverage was 85% for Basidiomycota, followed by Zygomycota with 93%. Primers were modified with 5’ sequencing adaptors (extended primer list in Online Resource 2), consisting of barcodes recommended by Roche and Lennon et al. (2010) and Lib-L adapters (Roche). PCR was conducted with AccuPrime Taq Polymerase High Fidelity (Invitrogen, USA) in a 40 µl reaction with the following conditions: initial denaturation for 3 min at 98°C followed by 32 cycles of 1 min denaturation at 94° C and 2 min annealing/elongation at 60°C. The quality of the amplicons were checked on a gel and we assured that the bands were of minor to moderate intensities, which is important for semi-quantitative considerations (Lindahl et al. 2013). PCR amplicons were purified using **AM**Pure XP Beads (Beckman Coulter) and quality was verified by microfluidics electrophoresis (Bioanalyzer, Agilent). The 9 **PCR** products per habitat (3 replicates per sampling time) were then pooled into three final replicates for sequencing, each of which contained included all 3 time points. As a result, the sequencing triplicates were representative for the habitat biota within the sampled timespan of April-June. We sequenced only one of the triplicates of the reed habitat. Afterwards all amplicons were pooled equimolar for emulsion PCR and subjected to pyrosequencing library preparation and sequencing following the manufacturer’s recommendations (FLX titanium chemistry, Roche). The sequence data was deposited at ENA (http://www.ebi.ac.uk/ena) under following accession number: PRJEB14236.

### Sequencing data processing

Raw 454 sequencing data were transformed by coding any nucleotide with a Phred score <11 as N. We removed all reads shorter than 300 nt and trimmed reads with trailing Ns. We then used two approaches for the biological classification of sequence reads: alignment-based clustering and alignment-free similarity. For the alignment-based clustering analysis, we generated OTUs using the distance-based approach in Mothur (Schloss et al. 2009). Filtered reads (fastq format) were first clipped using Shore (Ossowski et al. 2008). The D1 region is highly variable and has a pronounced length polymorphism, which renders an accurate alignment difficult. We therefore defined an end position to serve as an alignment anchor by screening the SILVA reference database (v123) for a conservative eukaryotic region located within our amplicon. We identified a conserved 42-nt sequence (GAGNCCGATAGNNNACAAGTANNGNGANNGAAAGWTGNAAAG) located after the D1 region as being suitable to serve as a stable 3′ end for the alignment. We manually designed a probe sequence in ARB (Ludwig et al., 2004) and subsequently clipped all reads after the last nucleotide. This normalized the length of the reads to a fixed position in a global alignment (average read length: 360 ± 13, n = 596k). We allowed for mismatches by scoring each match with 3, mismatches with -1, and gaps with -4. The threshold for clipping was set to score_MAx_>0.5 and the effect on the size-frequency distribution can be found in Online Resource 3. Unclipped sequences were rejected and analysed separately (Online Resource 3).

Clipped reads were processed in Mothur following 4-5-4 SOP (Schloss et al. 2009, accessed in August 2012). Quality filtering was achieved by using the sliding-window option (quality threshold of 25). For the alignment-based procedure, we constructed a reference dataset with long, high-quality reads processed with pyrotagger (http://pyrotagger.jgi-psf.org) using a cutoff at>500 nt. These were aligned to the eukaryotic backbone provided by the SILVA database LSURef (version 111; www.arb-silva.de) using the SINA aligner (Pruesse et al. 2012). This reference alignment was used to align our reads in Mothur. After clustering at 97% (average neighbour algorithm), the OTU abundance matrix was exported into R (www.r-project.org, version 3.3.1.) for further analysis (see below). Phylogenetic position of OTUs on the kingdom level was determined by adding the OTUs to the reference tree by the parsimony option implemented in ARB (Ludwig et al. 2004). As a comparison to a fixed 97% cutoff, we employed a coalescent-based clustering analysis as implemented by the gmyc model (Powell et al. 2011; Fujisawa and Barraclough 2013) for the early diverging lineages (541 OTUs, 3414 sequences).

The OTU sampling matrix (3682 OTUs, see Results) was subsampled at the lowest count of reads in a sample (1983 reads) to describe the overall alpha diversity measures for the habitats: Shannon index, community evenness, and a corrected Chao index (Chiu et al. 2014). Differences among habitats were examined with a non-metric multidimensional scaling (NMDS) ordination plot based on the Cao distance (Cao et al. 1997), which accounts for variable sampling intensity. Ellipses correspond to the standard-error around the habitat group centroids. The underlying clustering is based on UPGMA.

### Database-dependent sequence analysis

We performed two database-dependent analyses with the aim to resolve the taxonomic classifications of our sequences. First, clipped sequences were demultiplexed and quality trimmed in Mothur as described above and then submitted to SILVA NGS (www.arb-silva.de/ngs/) (Quast et al. 2013) for classification at the minimum similarity level of 85%. This resulted in 57 fungal taxonomic paths. Second, we performed an analysis without alignment, in which each replicate was pooled into sequences of one habitat, which were then subjected to a blast search (Blast+) against the nt database (Genbank) for all eukaryotes (Online Resource 4). Sequences were then classified using the LCA classifier implemented in Megan5 (Huson et al. 2011) using the following parameters: Min. Score=100, Max. Expected=0.01, Top %=5.0, Min. Support %=0.01, Min. Support=2, LCA=75%, Min. Complexity=0. Habitats were compared based on square root normalization.

## Results

### Diversity

Total eukaryotic alpha diversity estimated using alignment-based (97%) clustering was 3682 OTUs and the number varied considerably among habitats, with highest values found in Benthos and Biofilm habitats and lowest in Plankton and Reed habitats (Table 1). Of the total OTUs, 1042 (i.e. 28%) were fungi, according to their parsimony-based phylogenetic position (Table 1). The gmyc method of OTU delimitation for the non-Dikarya taxa (mainly aquatic lineages that comprised 52% of the fungal OTUs in our data) resulted in a 38% increase in the number of OTUs, suggesting that the 97% criterion underestimated taxon richness. However, the proportion of taxa that were unique to one habitat were similar in both analyses (97%: adj. r^2^=0.71; gmyc: adj. r^2^=0.74), thus we hereafter present OTUs based on the 97% criterion. The number of fungal OTUs per habitat was lowest in the Plankton (8 OTUs), higher in the water samples (Littoral, Pelagic, Above Sediment) and on the Reed (36-64 OTUs), and markedly higher in the Sediment, Biofilm, and Benthos samples (293-428 OTUs) (Table 1, Figure 2). Because the eukaryotic OTU count was positively correlated with read count (Table 1; Pearson’s *r*=0.90) we also estimated the total eukaryotic richness using the corrected Chao index. OTU richness was much higher using the Chao index, but habitats ranked similarly (lowest richness in Plankton; highest in Biofilm) (Table 1).

Fungal communities were clearly structured into different habitats according to the NMDS clustering of OTUs (97% criterion) (Figure 3). The three water samples (Pelagic, Littoral, and Above Sediment) and the Plankton sample all clustered together, whereas all other habitats were distinct (Figure 3). Very similar results were found by NMDS clustering of fungal taxonomic paths generated by SILVA NGS, indicating that the habitat clustering took place at even higher taxonomic levels (phylum to order level; data not shown).

### Taxonomic composition

Most sequences could only be classified as “fungi” or “environmental samples” based on the BLAST analysis using the nucleotide database of NCBI (mean: 70.2% of sequences; range: 36.5 %-89.7% of sequences in a given habitat) (see Online Resource 4). Thus, while BLAST may help to identify some individual sequences at a higher taxonomic resolution, we report the classification obtained from SILVA NGS for an overall comparison (Figure 4). OTUs could not be reliably classified to a taxonomic level lower than order because of database limitations. The orders Spizellomycetales and Rhizophydiales (both Chytridiomycetes) comprised the majority of sequences in the four pelagic habitats (Pelagic water, Above sediment water, Plankton, and Sediment) and the Littoral water sample, with a greater proportion of Spizellomycetales in the three types of water samples compared to more Rhizophydiales in the Plankton and Sediment habitats (Figure 4). In contrast, no single group was dominant in the Biofilm habitat (leading to the highest evenness in this habitat, Online Resource 1) with Chytridiales, Rhizophydiales (both Chytridiomycetes), and Agaricomycetes (Basidiomycota) forming the most prominent orders (Figure 4). Capnodiales and Helotiales (both Ascomycota) were the most prominent orders in the Benthic habitat whereas the Reed habitat was dominated by Pleosporales (Ascomycota) (Figure 4).

Only a small proportion of fungal sequences (0-6%) could be clearly assigned to forest taxa (Agaricales, Auriculariales, Boletales, Cantharellales, Gleophyllales, Hymenochaetales, Polyporales, Russulales; Online Resource 1). This value was significantly higher in the Sediment samples (mean=4.9 %, SD=0.9) than in all other habitats (ANOVA, F=11.36, p=0.000972; Tukey Post Hoc test, p<0.05; Reed, Littoral, and Plankton samples were excluded from the test due to the complete absence of these forest taxa). We recovered sequences from Oomycetes, a taxon that was formerly grouped with aquatic fungi and that occupies the same ecological niches (Sparrow 1960), in all samples (i.e. Albugo, Aphanomyces, Phytophthora, Pythium, Saprolegnia). These sequences were 1 - 3 orders of magnitude lower in abundance compared to the fungal sequences, with maximum values in the Benthos and Sediment samples (Online Resource 1).

**Figure 4.**
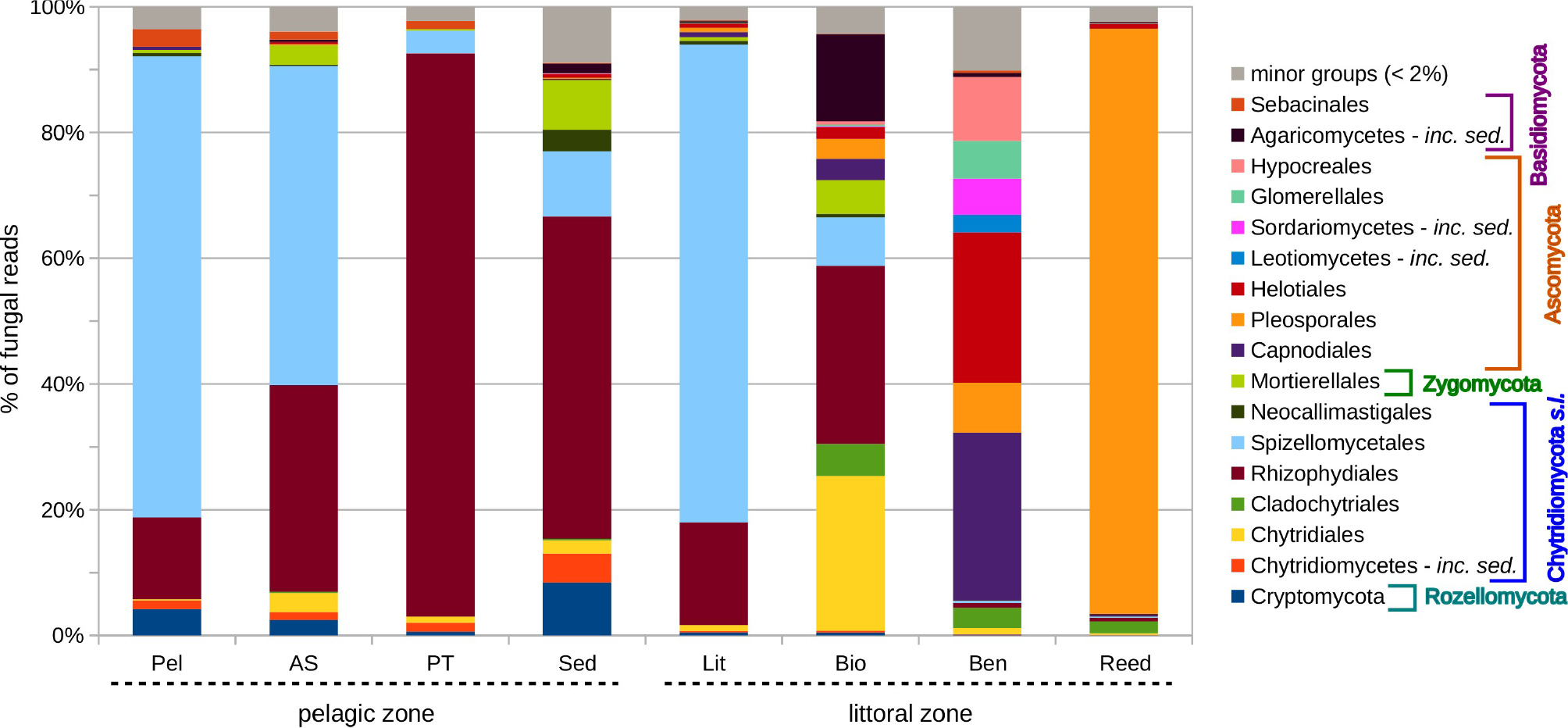
Distribution of fungal orders in Lake Stechlin habitats based on the mean percentages as determined with SILVA NGS. Fungal phyla are highlighted as brackets. Habitat codes are taken from Table 1.

## Discussion

Fungi play an important role in the cycling of carbon and nutrients in a wide range of freshwater habitats. While much of our understanding of their role in nutrient cycling stems from research in the terrestrial realm, there is increasing interest in their taxonomic and functional diversity in freshwater systems (Bärlocher and Boddy 2016; Grossart et al. 2016; Grossart and Rojas-Jimenez 2016). We assessed fungal diversity in eight different lake habitats, in contrast to most studies, which have compared water samples from different lakes (e.g., Monchy et al. 2011; Taib et al. 2013). We found marked differences in diversity and community composition among different habitat types in a single lake, and conclude that total fungal diversity was profoundly influenced by turnover in the taxonomic composition among different habitats. A highly exciting result was the finding of high fungal diversity in biofilms, suggesting that for such habitats future studies need to simultaneously target both recently and early diverging groups (e.g., Dikarya and Rozellomycota, respectively). As predicted, we observed a change in fungal phyla from open-water habitats compared with those that were more dominated by CPOM. In the following discussion, we first address methodological considerations in the study and then discuss fungal diversity separately for the major habitats in order to evaluate habitat-specificity of the fungal community structure in detail.

### Methodological considerations

The occurrence of both, evolutionary distant early diverging fungal lineages and members of the Dikarya renders a comprehensive assessment of the aquatic mycobiota challenging and requires a careful choice of the DNA marker. In this study, we employed the D1 region of the LSU as a barcode because of its high variability while still being conservative enough to amplify across the fungal kingdom. Both D1 and D2 regions were formerly used as a molecular marker for fungi, especially for yeast identification (Kurtzman and Robnett 1997) and perform almost as well as the commonly used ITS region in discriminating fungal groups (Schoch et al. 2012). Moreover, the LSU is an established phylogenetic marker for Chytridiomycota (Letcher et al. 2006) and unlike the ITS region it can be used to delimit distant aquatic fungal lineages (Lefèvre et al. 2012; Wurzbacher et al. 2014). The small ribosomal subunit (SSU) is also well established for early diverging lineages (e.g., Jobard et al. 2012; Ishii et al. 2015), however, it is less suitable for fungal groups within Dikarya (Tedersoo et al. 2015) and would fall short for complex habitats such as our biofilms. Currently, the major drawback of using LSU regions as a phylogenetic marker is the lack of a suitable database. Until now, there exists no well curated LSU database for fungi. The most recent curated SILVA LSU (v123) database just contains 1038 full-length sequences. It may be possible to use the truncated/partial datasets of SILVA and RDP (Fungal 28S rRNA Sequences dataset) which are constantly updated and contain more than 100k fungal entries for the D1 region, but this may also increase the risk of false annotations, since aquatic fungi are largely underrepresented (Panzer et al. 2015).

### Water and large plankton

All three water habitats had a low proportion of fungal sequences, but had almost an order of magnitude higher proportion of total OTU diversity. We observed a predominance of Chytridiomycota in water samples and in the habitats directly connected to processes in the open water, namely large planktonic organisms (PT) and Sediment. The proportion of fungal sequences in these samples was relatively low (see also Luo et al. (2011) for previous results on Lake Stechlin) and this may relate to the fact that we did not enrich for fungi by prefiltration or by primer selection. Lefèvre et al. (2012) provided a summary of fungal and chytrid percentages ranging from 1 - 50% in water samples and how this may relate to prefiltration and the employed primer pair. Like Monchy et al. (2011), we observed similar communities in the water samples (Littoral, Pelagic, and Above Sediment). In pelagic water, we found an increased proportion of Rozellomycota and the deep water also had increased proportions of Mortierellales. It might be that parasitic fungal groups, in particular, are part of the rare plankton community (Mangot et al. 2013), whereby they can recruit for temporal infection events on time scales of a few weeks as has been observed in many studies (e.g., Ibelings et al. 2004; Alster and Zohary 2007). Between infections, their abundance may remain low, explaining the higher proportion of fungal taxa compared to fungal sequences.

There was a limited number of fungal taxa associated with zoo-and phytoplankton samples (PT; > 55 µm), which presumably should represent attached or infective stages of fungi. 84-93% of these fungi belonged to Rhizophydiales, a group of well described phytoplankton parasites. Interestingly, in the pelagic water samples Rhizophydiales only accounted for 14-17%. This is insofar important, because most microscopic studies on chytrids refer to infected algae of approximate the size of 50 µm or larger (e.g., Hohlfeld 1998; Ibelings et al. 2004; Rasconi et al. 2012). However, in an unbiased water sample (i.e. not fractionated by filtration or enriched by a plankton net), they were replaced as dominant group by the order Spizellomycetales, which are common saproptrophs in soil and may underline to the importance of saprotrophic chytrids in aquatic environments (Wurzbacher et al. 2014). This may establish the mycoloop (Kagami et al. 2014) as a trophic link during times with low prevalences of algal infections. Finally, there was a minute amount of the enigmatic Rozellomycota in the large plankton. Rozellomycota are discussed as *inter alia* attached algal parasites (Jones et al. 2011); however, our results indicate that they were not parasites of the larger plankton in Lake Stechlin, where Chytridiomycota fully occupied this niche. Due to the small size of Rozellomycota, they may rather have a specialization towards smaller hosts, which do not provide enough resources for Chytridiomycota to complete their life cycle.

### Sediment

The profundal surface sediments of Lake Stechlin were rather similar to the water sample in terms of fungal community composition. The profundal sediment temperature in Lake Stechlin remains around 4°C and the upper sediment surface (~ 5 mm) is usually oxic. The sediment has a high water content (> 95%) and high organic matter content at the sampled sites because it receives mainly sinking matter from pelagic organisms. Thus the sediments serve as an organismic archive and it was therefore not surprising to find slightly elevated proportions of forest fungi that were probably washed in as spores. The dominant fungal group was the Rhizophydiales, similarly to the large plankton samples. Like their hosts, parasitic chytrids develop thick-walled resting spores (cysts), which can be found in the sediment, and there are parasitic chytrids that infect algal resting stages in sediments (Canter 1948, Canter 1968). The sediments also had increased proportions of Rozellomycota, Mortierellales, and Neocallimastigomycota. The few studies that have investigated lake or pond sediments reported Chytridiomycota and Rozellomycota (at that time referred to as LKM11 & LKM15) to be the dominant fungal phyla (Luo et al. 2005; Slapeta et al. 2005). Rozellomycota appear to occur in the hypolimnion of lakes (Lepère et al. 2010) and can also be found in anoxic habitats (Jones et al. 2011); however, their ecological function remains unclear (Grossart et al. 2016). Similarly enigmatic was the appearance of Zygomycota at the sediment surface. They are most likely saprotrophs and they are known to grow at low temperatures under oxic conditions, e.g. under snow packs in sub-alpine regions (Schmidt et al. 2008). Very surprising was the appearance of Neocallimastigomycota, which are by definition obligate, mutualistic, anaerobic rumen fungi. They are exceptional in that they break down a broad variety of plant polymers under anaerobic conditions (Solomon et al. 2016). These fungi must have had an environmental ancestor and it is possible that anoxic sediments may provide a suitable habitat for these anaerobic fungi. However, the sequences were only approx. 90% similar to *Orpinomyces,* and it is likely that they constitute a sister clade. Lefèvre et al. (2012) also described sequences from lake plankton samples that may support such a new environmental lineage of “rumen fungi” and it remains open whether or not those sequences cluster phylogenetically inside the Neocallimastigomycota.

### Biofilm (Periphyton)

Biofilm samples appeared to represent an intermediate fungal habitat between sediments and benthic samples by including a high diversity of early diverging lineages and elevated proportions of Dikarya (16 - 36%, encompassing similar proportions of Basidiomycota (18%) and Ascomycota (12%)). Fungi seem to represent a significant proportion of the overall eukaryotic biofilm community, dominated by biofilm-forming algae (the ratio of fungi to periphyton/epilithic algae was roughly 1:5). Biofilms represent a complex environment (exhibiting the highest eukaryotic taxon richness of all eight habitats) and this is also reflected by a high diversity of fungal groups and taxa. Next to Rhizophydiales and Spizellomycetales, we also found high proportions of other chytrids of the orders Chytridiales and Cladochytridiales. These autotrophic lake biofilms seem to be a rich source of fungal biodiversity and pose a very interesting habitat for future studies. Biofilms (in our case mainly littoral periphyton and epilithic biofilms) have been rarely examined for fungi, and only a few studies on stream ecosystems have investigated the fungal occurrence (measured as ergosterol) on substrates other than leaves (Tank and Dodds 2003; Artigas et al. 2004, Aguilera et al. 2007; Frossard et al. 2012). In lakes and streams, periphyton can contribute substantially to the primary production of the whole ecosystem (Lalonde et al. 1991; Vadeboncoeur et al. 2007 and references therein; Vis et al. 2007), and represents the main food source for macrozoobenthic grazers (Cattaneo and Mousseau 1995). It is a habitat of high ecological relevance and our findings suggest that it is not only a rich source of fungal biodiversity, but that fungi might play an important ecological role in this habitats, turning over a significant amount of the “living” algal carbon, and thus total carbon in the lake.

### Benthic and reed samples (CPOM)

In contrast to water samples, fungal sequences were the dominant organism when the substrate was CPOM. Fungi were the dominant biotic component in reed and benthic samples, which mainly consisted of submerged plant residues (in addition to certain algae and benthic animals). Mitosporic ascomycetes lineages were predominant, followed by a small percentage of chytrids (mainly Cladochytridiales) and almost no Basidiomycota sequences. Mitosporic ascomycetes are superior plant decomposers in freshwater systems (Gessner et al. 2007), where they are known as aquatic hyphomycetes. We could not clearly assign our sequences to the species level of known aquatic hyphomycetes because of a lack of reference sequences in public databases (see discussion below) and further investigations will have to confirm a match. The Benthos habitat had a high diversity of fungal OTUs and is probably home to those fungi responsible for the breakdown of submerged plant remains. Interestingly, the importance of aquatic hyphomycetes for plant litter breakdown has thus far only been demonstrated in lotic environments, and lakes have not been investigated in detail (see Chauvet et al. 2016). In contrast to the benthic samples, the Reed sample exhibited a very restricted diversity of fungi, mainly related to the fungal order of Pleosporales (93*%).* Early molecular work has already established the high diversity of reed endophytes (Neubert et al. 2005) and we could confirm their presence, although our diversity estimates are lower than the estimates of Neubert et al. (2005). The reed sample can be seen as an outgroup in our study, also since it comprised emergent parts of plants. Sequences of the order Pleosporales were largely restricted to the reed and to a small extent to benthic samples.

## Conclusions

Our broad screening revealed a very high and habitat-specific biodiversity of fungi in a single lake. It extends previous research in freshwaters and clearly indicates that the sediment and various biofilms are hotspots of aquatic fungal diversity. Most of the recovered fungi can be considered as indigenous and only a very small proportion of fungi originated from the terrestrial surroundings, except for sediments which may presumably also collect fungal spore from the terrestrial environment. Our study aims to stimulate further research on the thus far undersampled lake habitats, such as various sediments and biofilms, sediments in the euphotic zone, and biofilms on submerged macrophytes (Kohout et al. 2012). A more holistic approach in evaluating fungal diversity should also enable a deeper insight into the multifunctional ecological roles of fungi in limnetic ecosystems with differing environmental features. This is of particular importance to evaluate the relationship between biodiversity and ecosystem function.

## Acknowledgments

Research was partially supported by the Leibniz SAW/Pakt for Research and Innovation project “MycoLink” and by an IGB Fellowship in Freshwater Science awarded to NW, HPG, and MTM. CW acknowledges a Marie Skłodowska-Curie post doc grant (660122, CRYPTRANS). This is publication 39 of the Berlin Center for Genomics in Biodiversity Research.

